# Biophysical insight into protein-protein interactions in the Interleukin-11/Interleukin-11Rα/glycoprotein 130 signaling complex

**DOI:** 10.1101/2022.07.14.499993

**Authors:** Chinatsu Mori, Satoru Nagatoishi, Ryo Matsunaga, Daisuke Kuroda, Makoto Nakakido, Kouhei Tsumoto

## Abstract

Interleukin-11 (IL-11) is a member of the interleukin-6 (IL-6) family of cytokines. IL-11 is a regulator of multiple events in hematopoiesis, and IL-11-mediated signaling is implicated in inflammatory disease, cancer, and fibrosis. All IL-6 family cytokines signal through the signal-transducing receptor, glycoprotein 130 (gp130), but these cytokines have distinct as well as overlapping biological functions. To understand IL-11 signaling at the molecular level, we performed a comprehensive interaction analysis of the IL-11 signaling complex, comparing it with the IL-6 complex, one of the best-characterized cytokine complexes. Our thermodynamic analysis revealed a clear difference between IL-11 and IL-6. Surface plasmon resonance analysis showed that the interaction between IL-11 and IL-11 receptor α (IL-11Rα) is entropy driven, whereas that between IL-6 and IL-6 receptor α (IL-6Rα) is enthalpy driven. Our analysis using isothermal titration calorimetry revealed that the binding of gp130 to the IL-11/IL-11Rα complex results in entropy loss, but that the interaction of gp130 with the IL-6/IL-6Rα complex results in entropy gain. Our hydrogen-deuterium exchange mass spectrometry experiments suggested that the D2 domain of gp130 was not involved in n IL-6-like interactions in the IL-11/IL-11Rα complex. It has been reported that IL-6 interaction with gp130 in the signaling complex was characterized through the hydrophobic interface located in its D2 domain of gp130. Our findings suggest that unique interactions of the IL-11 signaling complex with gp130 are responsible for the distinct biological activities of IL-11 compared to IL-6.

## Introduction

The interleukin-6 family cytokines, including interleukin-6 (IL-6), interleukin-11 (IL-11), leukemia inhibitory factor (LIF), and oncostatin M (OSM), regulate complex cellular processes such as gene activation, proliferation, and differentiation (1). Although all IL-6 family proteins transduce signals through a common receptor, glycoprotein 130 (gp130), they have distinct as well as overlapping biological functions (2–6). The mechanisms that underlie these functional redundancies and distinctions have not been elucidated (2, 7).

IL-11 is as a key regulator of multiple events in hematopoiesis (8). As IL-11 can promote maturation of platelets (9), early research on IL-11 focused on it role in thrombocytopenia (10–12). More recent studies have shown that IL-11 can stimulate fibrosis and tumor progression by inhibiting apoptosis and promoting the proliferation of cancer cells (13–15) and that it is necessary for organ regeneration in tadpoles (16). Because IL-11 relates to such various diseases and has complex and elusive biological activities, it is focused on as an important subject of research.

The interaction between IL-11 and its receptor IL-11 receptor α (IL-11Rα) has been characterized using biophysical techniques (17), and in silico analysis of the interaction among IL-11, IL-11Rα, and gp130 has been reported (18). Molecular level analyses have been hampered by difficulties in the preparation of recombinant IL-11 and IL-11Rα. The complex of IL-6 and IL-6 receptor α (IL-6Rα) has been well characterized. Both IL-11/IL-11Rα and IL-6/IL-6Rα bind to gp130, to form heterotetramers (19, 20).

In this report, we describe our comprehensive biophysical interaction analysis of the IL-11/IL-11Rα/gp130 signaling complex using surface plasmon resonance (SPR), isothermal titration calorimetry (ITC), and hydrogen/deuterium exchange mass spectrometry (HDX-MS), and molecular modeling using ColabFold (21–23). We compared our findings to similar experiments performed with the IL-6 complex. Characterized IL-6 family cytokine signaling complexes interact with gp130 through the hydrophobic interface located in its D2 domain (24–26). However, our data show that IL-11/LI-11Rα does not interact with the D2 domain of the gp130 complex. This unique interaction mode may underlie the distinct biological activities of IL-11 and suggest a target for IL-11-specific therapeutic development.

## Results

### Surface plasmon resonance analysis of the interaction between interleukins and their receptors

To evaluate the thermodynamics of the interactions between IL-11 and IL-11Rα, a stable recombinant IL-11Rα was prepared using Mimic™ Sf9 cells. IL-11Rα was expressed and purified as a fusion protein with the Fc domain of human IgG1 (Figure S1) as reported previously (27, 28). IL-11 was prepared using an *Escherichia coli* expression system (Figure S2). Both IL-6 and IL-6Rα were expressed using Expi293F™ cells (Figure S3, S4).

We performed a van’t Hoff analysis of SPR data to evaluate the thermodynamic parameters of the interactions between IL-11 and IL-11Rα and between IL-6 and IL-6Rα (Table S1 and S2, Figure S5). For the interaction between IL-11 and IL-11Rα, Δ*H*, –*T*Δ*S*, and Δ*G* were 47±3 kJ/mol, -100±3 kJ/mol, and -53±4 kJ/mol, respectively (Figure 1, Table S3). Thus, the interaction between IL-11 and IL-11Rα is largely entropy driven. These results are in agreement with a previous report (17). For the interaction between IL-6 and IL-6Rα, Δ*H*, –*T*Δ*S*, and Δ*G* were –49±5 kJ/mol, 6±5 kJ/mol, and –43±7 kJ/mol, respectively (Figure 1, Table S3). Therefore, the binding of IL-6 to IL-6Rα is enthalpy driven.

**Figure 1.**
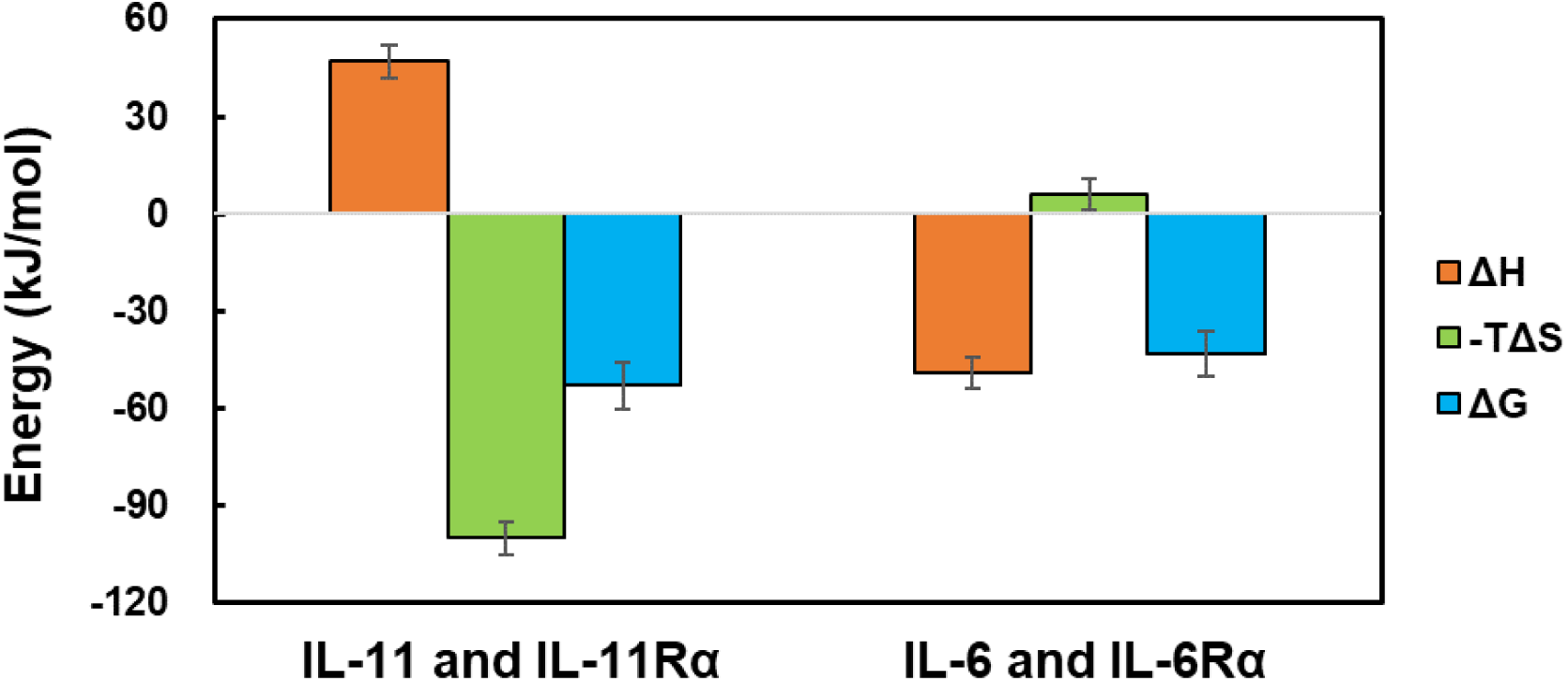
Thermodynamic parameters of the interaction between IL-11 and IL-11Rα (left) and between IL-6 and IL-6Rα (right) calculated from van’t Hoff plots of SPR data.

### Thermodynamics of the interaction between gp130 and the interleukin/receptor complexes

To examine the thermodynamics of the interaction between gp130 and the complexes between IL-11 and IL-6 and their receptors, recombinant IL-11/IL-11Rα and IL-6/IL-6Rα complexes were prepared. In these complexes, the interleukin and the receptor were connected with a glycine-serine linker (GGGGS)_3_, and the fused proteins were expressed in Expi293F™ cells. Hydrogen-deuterium exchange mass spectrometry (HDX-MS) analysis was performed to determine whether the interleukin and the receptor interacted properly in the fused forms. The deuterium ratios decreased at positions expected based on previous data (19, 29) to be located in the interaction interfaces of both the IL-11/IL-11Rα and IL-6/IL-6Rα constructs (Figure S6).

We evaluated the thermodynamics of the interaction between gp130 and recombinant complexes using ITC (Figure 2A, B). The gp130 interaction with IL-11/IL-11Rα was associated with the enthalpy gain (Δ*H* = –51±1 kJ/mol) and entropy loss (–*T*Δ*S* = 5±1 kJ/mol) (Figure 2C, Table S4). In contrast, the interaction of gp130 with IL-6/IL-6Rα resulted in gains in both enthalpy and entropy (Δ*H* = –28±1 kJ/mol, –*T*Δ*S* = –13±2 kJ/mol) (Figure 2B, Table S4). Thus, there were considerable differences in the thermodynamic parameters of the interactions not only between the interleukins and their receptors, but between gp130 and the interleukin/receptor complexes.

**Figure 2.**
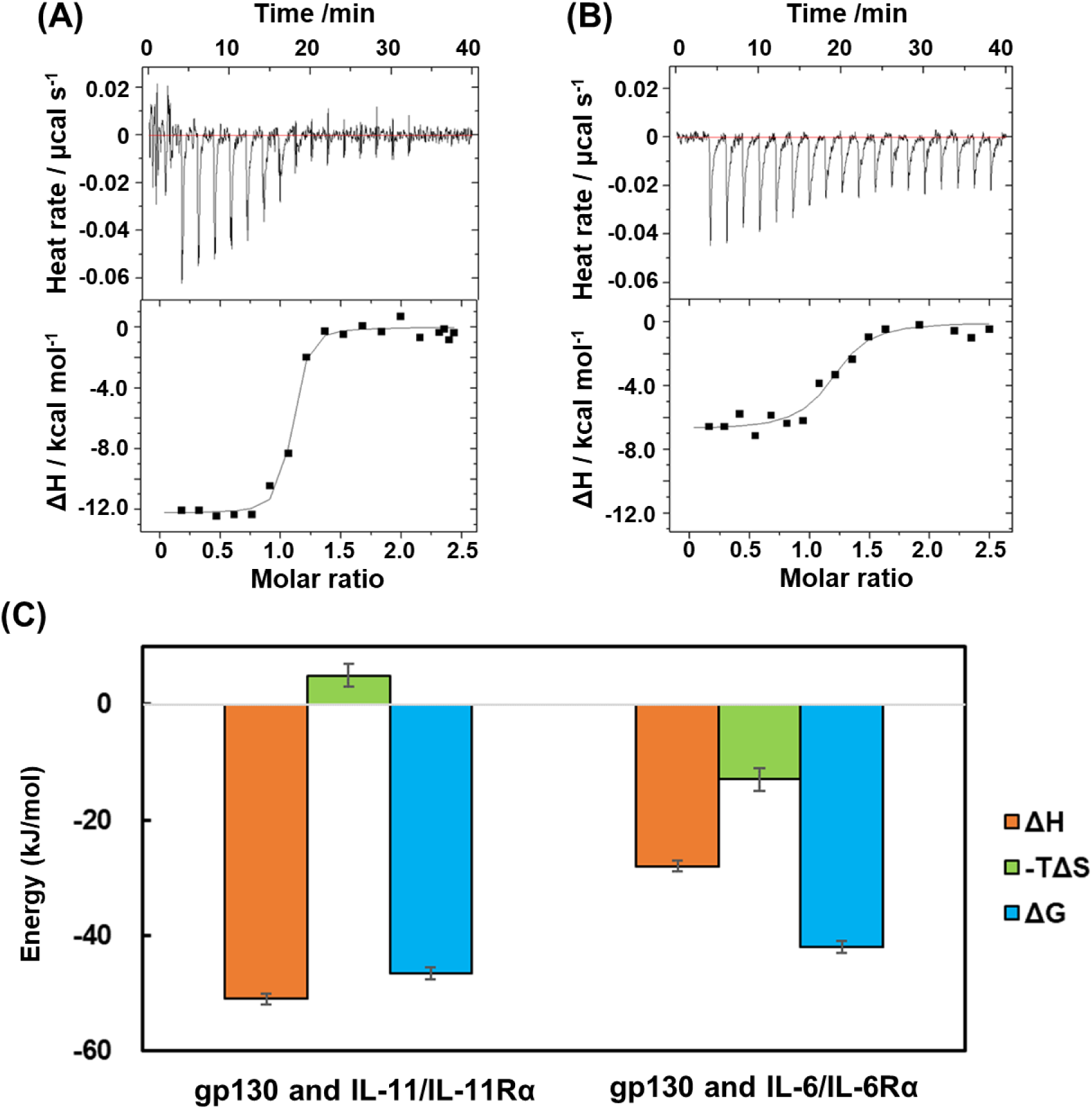
Thermodynamic analysis of gp130 interactions with interleukin/receptor complexes. (A) The ITC profile of the interaction of gp130 with IL-11/IL-11Rα. (B) The ITC profile of the interaction of gp130 with IL-6/IL-6Rα. (C) The thermodynamic parameters of the interactions between gp130 and interleukin/receptor complexes.

### Structural analysis of the interaction between gp130 and interleukin/receptor complexes

To obtain the information regarding the positions of the binding interfaces and the conformational changes that occur when gp130 interacts with interleukin/receptor complexes, we performed HDX-MS experiments. HDX rates of differences between gp130 alone and gp130 in complex with the interleukin/receptor fusions showed differences in HDX signals (Figure 3A-B). We assume that the IL-11/IL-11Rα/gp130 complex is also hexameric with a 2:2:2 stoichiometry, as is the IL-6 complex. For the interaction with IL-11/IL-11Rα, the differences in HDX rates between gp130 alone and in the complex involve amino acids 89-96 in the D1 domain and 254-261 in the D3 domain of gp130 (Figure 3C-F). Differences in HDX rates were also observed in these regions between gp130 alone and gp130 in the complex with IL-6/IL-6Rα as well as in the regions of amino acids 30-35 in the N-terminal region of the D1 domain and in the region of amino acids 160-166, which is the EF-loop in the D2 domain (Figure 3C-F).

**Figure 3.**
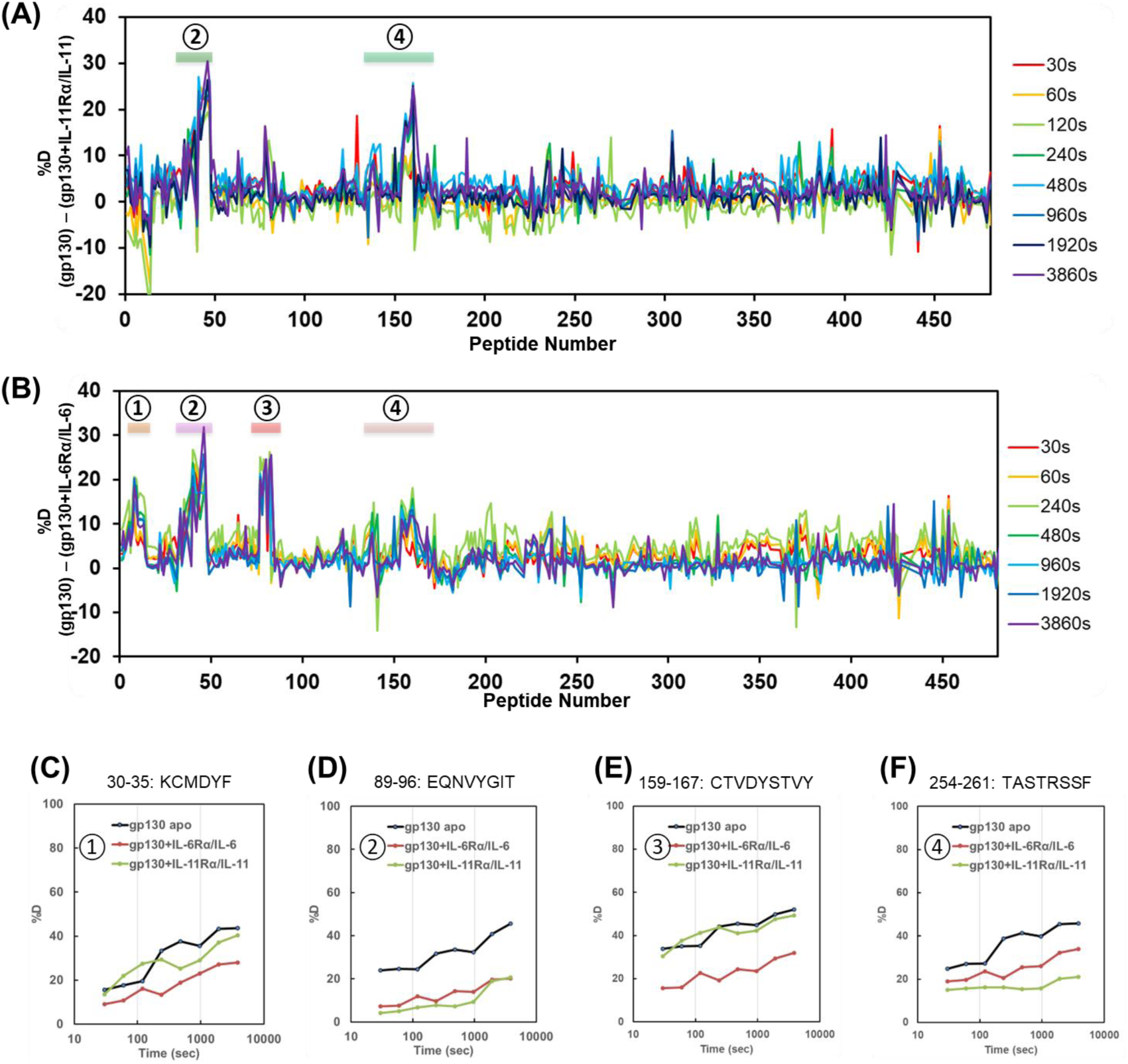
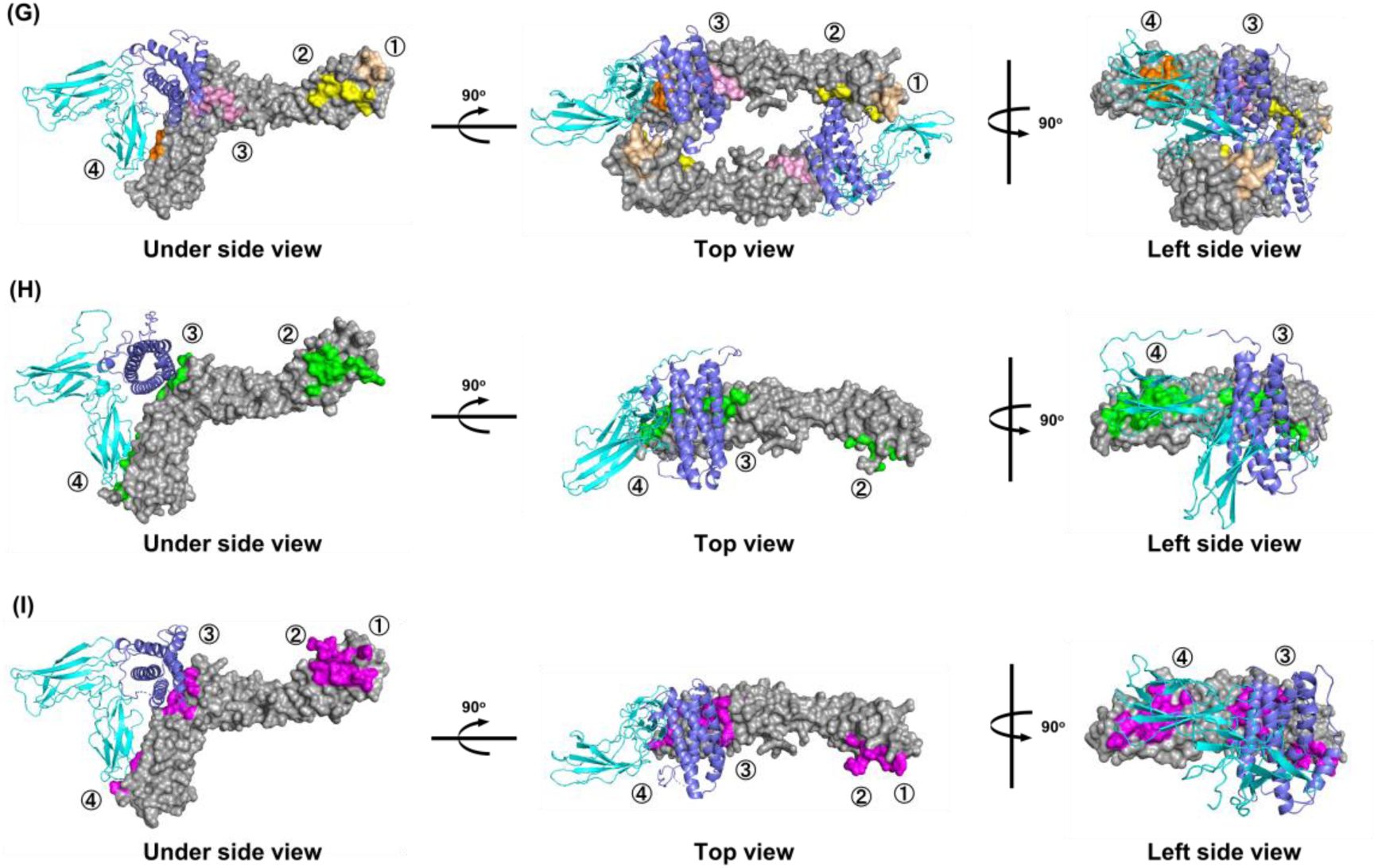
Interface analysis of gp130 with interleukin/receptor complexes. (A) Differences between HDX rates for gp130 in the absence and presence of IL-11/ IL-11Rα complex. (B) Differences between HDX rates for gp130 in the absence and presence of IL-6/IL-6Rα complex. (C-F) HDX-MS profiles of gp130 amino acids C) 30-35, D) 89-96, E) 159-167, and F) 254-261. (G) The predicted structure of the IL-6 signaling complex. gp130 is shown in gray with color indicating where HDX rates decreased. IL-6 is shown in light blue and IL-6Rα in dark blue. (H) The predicted structure of the IL-11 signaling complex. gp130 is shown in gray with green indicating amino acids buried more than 60% by the complex formation. IL-11 is shown in light blue and IL-11Rα in dark blue. (I) The predicted structure of the IL-6 signaling complex. gp130 is shown in gray with magenta indicating amino acids buried more than 60% by the complex formation. IL-6 is shown in light blue and IL-6Rα in dark blue.

A model of the IL-11 signaling complex was built using ColabFold (21–23). The crystal structure of the IL-6 signaling complex is available (19). To determine the contributions of each amino acid residue to the interaction at the interaction interface, we calculated the buried surface areas using the PISA server (30) (Figure 3G-I). For D1 (41-98) and D3 domains of gp130, the buried surface areas involving D1 (41-98) (733.06 Å^2^) and D3 (872.84 Å^2^) domains in gp130 in the IL-11/IL-11Rα/gp130 complex were higher than those of D1 (41-98) (494.34 Å^2^) and D3 (627.43 Å^2^) domains in the IL-6/IL-6Rα/gp130 complex (Table 1). Analysis of the contact sites between amino acids in the predicted and known structures of the signaling complexes suggested the sites of amino acid contacts in the D1 (41-98) and D3 domains were consistent with the amino acid sequence regions obtained in the HDX-MS analysis for both IL11 and IL6. (Figure S7).

**Table 1.**
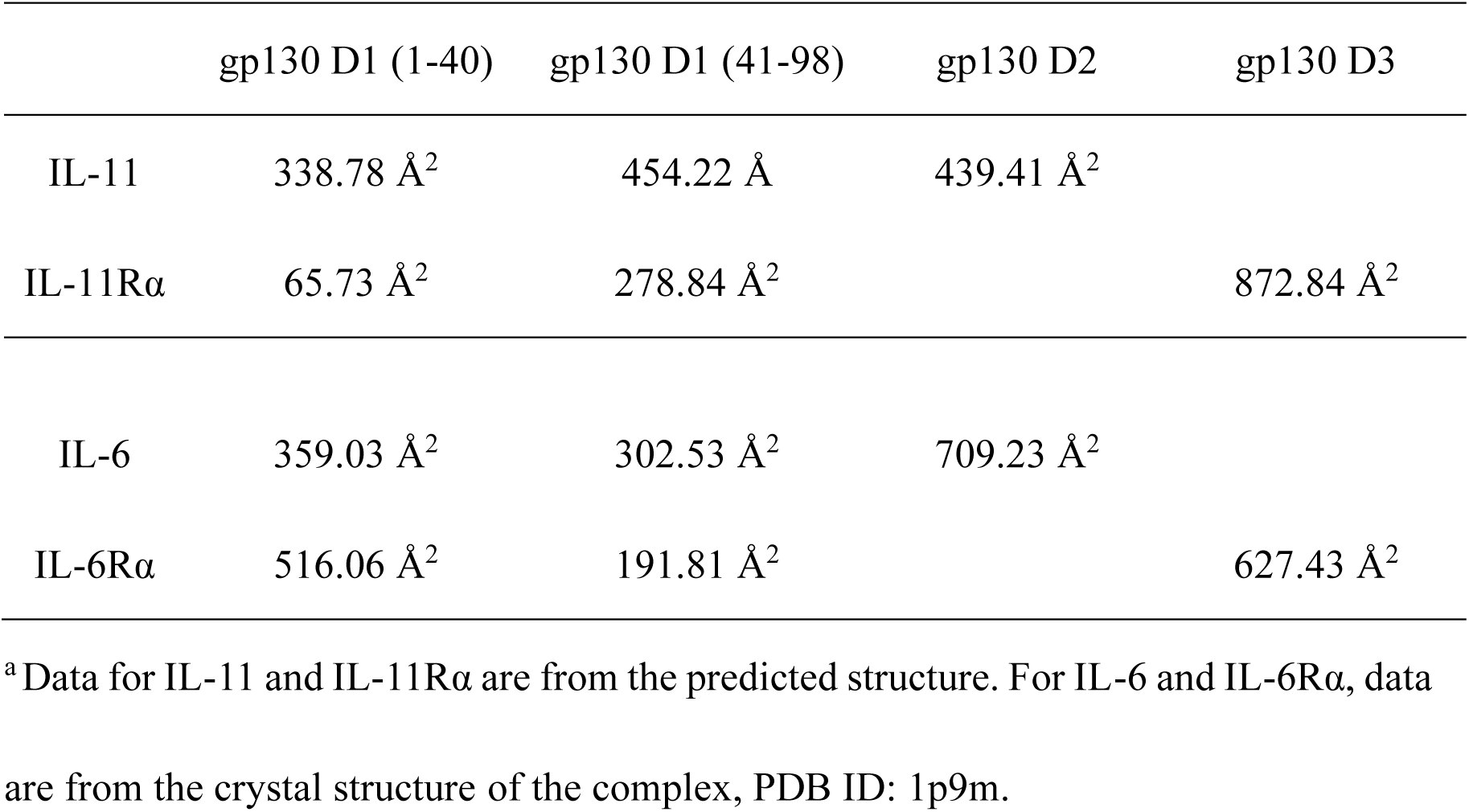
Buried surface areas of gp130 in IL/ILRα/gp130 complex^a^.

For the D1 (1-40) and D2 domains of gp130, the buried surface areas involving D1 (1-40) (404.51 Å^2^) and D2 (439.41 Å^2^) domains in gp130 in the IL-11/IL-11Rα/gp130 complex were lower than those of D1 (1-40) (875.09 Å^2^) and D2 (709.23 Å^2^) domains in the IL-6/IL-6Rα/gp130 complex (Table 1). These results are consistent with the finding that there were no differences in HDX rates were observed in the region of 30-35 in D1 and the region of 160-166 in D2 for gp130 in complex with IL-11/IL-11Rα compared to gp130 alone. The EF-loop of the D2 domain in gp130, amino acids 159-167, was not part of the binding interface with IL-11, whereas it was buried in the complex with the IL-6/IL-6Rα construct (Figure 3G-I). Analyses of the predicted and known structures of the signaling complexes suggested that hydrophilic amino acids were more involved in the interaction of the D2 with IL-11, whereas hydrophobic amino acids were involved in this interaction with IL-6 (Table S5), as reported previously (24–26). This difference may be the reason that the interaction of gp130 with IL-11/IL-11Rα is enthalpy driven, whereas that with IL-6/IL-6Rα is entropy driven.

Our HDX-MS data also suggest that the IL-11 alone forms a more rigid structure than does IL-6 alone (Figure S8A, B). In the complex with its receptor, the flexibility of IL-6 is not significantly different from that of the IL-6 apo form; in contrast, IL-11 is more rigid in the complex (Figure S8C, D).

## Discussion

Both IL-6 and IL-11 are four-helical bundle cytokines with an up-up-down-down topology, and both signal through gp130 (19, 20, 31–33). The two proteins have unique functions, and the goal of this study was to determine the molecular level interactions that lead to these functional differences. The interactions between the interleukins and their receptors were examined by SPR experiments. The SPR data showed that the interaction between IL-11 and IL-11Rα was entropy driven, resulting from the previously described hydrophobic interface between IL-11 and IL-11Rα (17). Our thermodynamic parameters differ from those of Metcalfe et al. (17) due to the difference in the measurement system. In our experiments, the receptor was immobilized for SPR, whereas Metcalfe et al. evaluated binding in solution using ITC. We infer that immobilized IL-11Rα has a lower loss of structural entropy and that our thermodynamic parameters have more favorable entropy values than those determined by ITC.

We further examined the interactions between gp130 and IL-11/IL-11Rα and between gp130 and IL-6/IL-6Rα using ITC experiments and HDX-MS. Our ITC experiments revealed that the interaction of gp130 with IL-11/IL-11Rα was associated with entropy loss, whereas the interaction of gp130 with IL-6/IL-6Rα was associated with entropy gain. Previous work established that the interactions of gp130 with IL-6 family cytokines IL-6, LIF, and OSM are associated with entropy gain (24, 34, 35), so IL-11 is unique among this family. Moreover, HDX-MS analyses showed that there are differences in the binding interfaces of gp130 on IL-11/IL-11Rα versus IL-6/IL-6Rα. The D2 domain of gp130 recognizes the IL-6 family cytokine complexes through hydrophobic interactions (24–26). Our data showed that the EF-loop of the D2 domain in gp130 was not part of the binding interface with IL-11/IL-11Rα, whereas it was buried in the complex with the IL-6/IL-6Rα construct. Our experiments do not rule out the possibility that a few residues located in the D2 domain of gp130 recognize IL-11. In fact, previous mutational studies revealed that Phe 191, which is located in the D2 domain, is crucial for binding to both IL-6 and IL-11 (36–38). The fact that IL-11 forms the signaling complex without hydrophobic interactions with the D2 domain could explain entropy loss upon binding that was revealed by the ITC experiment. HDX-MS analysis also revealed that IL-11 was more rigid than IL-6 in the complex with the respective receptor. This difference in the structural flexibility between IL-11 and IL-6 may impact interactions with gp130.

Previous studies have provided insight into the structure of the IL-6 signaling complex (39), and recent work had suggested differences between the IL-11/IL-11Rα/gp130 complex and the complex of gp130/IL-6/IL-6Rα (7, 17, 40). Our study provides the first biophysical evidence for differences in the interactions of these cytokines with gp130. We hypothesize that these differences influence the behavior of the intracellular domain of gp130 and differences in signal intensity or changes in the balance of preferentially activated signaling pathways (e.g., JAK/STAT pathway versus the Raf/ERK pathway). Our study will guide further analyses such as mutation studies that will reveal in more detailed the molecular mechanisms of IL-11-mediated signaling complex, which will in turn lead to development of cytokine-specific drugs.

### Experimental procedures

#### Expression and purification of proteins

Information of the constructs of IL-11, IL-11R*α*, IL-6, and IL6R*α*, and their expression and purification are shown in the supporting information. In constructs of interleukin/receptor complexes, the interleukin and the receptor were connected with a glycine-serine linker (GGGGS)_3_. The IL-11/IL-11Rα fusion contained amino acids 113-318 of IL-11Rα and amino acids 23-199 of IL-11, and the IL-6/IL-6Rα fusion contained amino acids 114-320 of IL-6Rα and amino acids 30-212 of IL-6. The sequence for human gp130 (amino acids 24-618) containing a myc-tag and a hexahistidine-tag at the C-terminus was subcloned into plasmid pcDNATM 3.4-TOPO® vector (Thermo Fisher Scientific). For the expression of IL-6, IL-6Rα, and gp130, the Expi293™ Expression System (Thermo Fisher Scientific) was used. The transfection of Expi293F™ cells was performed according to the manufacturer’s protocol. Enhancer 1 and Enhancer 2 were added to transfected cultures at 20 h post-transfection. The cells were incubated at 37 °C, 125 rpm, 7% CO2 for 3 days. Proteins were purified from the supernatant using Ni-NTA agarose (Qiagen) at 4 °C. Bound proteins were eluted with 20 mM Tris-HCl (pH 8.0) 500 mM NaCl 500 mM imidazole. Final purifications were performed by size exclusion chromatography (SEC) on HiLoad^®^ 16/600 Superdex^®^ 200 pg (GE Healthcare) with running buffer composed of 20 mM HEPES-NaOH (pH 7.4) and 150 mM NaCl.

### Kinetics analyses using surface plasmon resonance

The interactions between interleukins and their receptors were analyzed by SPR with a Biacore T200 instrument (GE Healthcare). To immobilize the receptors, a human antibody capture kit (GE Healthcare) was used. Anti-human Fc IgG was immobilized on reference and sample flow cells of Series S CM5 sensor chip (GE Healthcare) by amine coupling according to the manufacturer’s protocol. IL-6Rα or IL-11Rα was then captured. The kinetic assay was carried out in 20 mM HEPES-NaOH (pH 7.4), 150 mM NaCl, 0.005% (v/v) Tween 20 containing IL-11 at concentrations from 0.41 nM to 33 nM or IL-6 at concentrations from 13 nM to 200 nM at a flow rate of 30 µL min^-1^. The binding curves were obtained by subtracting the binding response on the reference flow cells from that on the sample flow cells. Kinetic parameters were calculated by global fitting analysis assuming a Langmuir binding model and a stoichiometry of 1:1. The dissociation constant (*K*_*D*_) was determined from the equation *K*_*D*_ = *k*_*off*_ / *k*_*on*_. A van’t Hoff thermodynamic analysis of the interaction was carried out by repeating SPR measurement at different temperatures. The thermodynamic parameters were calculated from plots of ln *K*_*D*_ versus 1*/T*.

### Isothermal titration calorimetry analyses

Calorimetric titration analyses were carried out on ITC200 calorimeter (MicroCal) at 25 °C. All experiments were carried out in 20 mM HEPES-NaOH (pH 7.4), 150 mM NaCl. gp130 (35 µM or 42 µM) was put in the syringe, and 2.4 µM IL-11/IL-11Rα or 4.2 µM IL-6/ IL-6Rα was put in the cell. Data were processed with the MicroCal Origin 5.0 software.

### Molecular modeling of IL-11/IL-11Rα/gp130 complex and identification of interface residues

A model of the IL-11/IL-11Rα/gp130 complex structure was made using Google AlphaFold2_mmseq2 notebook from the ColabFold project (21–23). IL-11 and IL-11Rα were connected with a (GGGGS)_3_ linker. The interface residues of the model structure of the IL-11/IL-11Rα/gp130 complex and of the crystal structure of the IL-6/IL-6Rα/gp130 complex were defined using the PDBe PISA server (30).

### Hydrogen-deuterium-exchange mass spectrometry

Each protein sample was prepared in 20 mM HEPES-NaOH (pH 7.4) and 150 mM NaCl at a final concentration of 1.0 mg mL^-1^. The protein solution was diluted 10-fold with the same buffer prepared in D_2_O. The diluted solutions were then incubated separately at 10 °C. Deuterium-labeled samples were quenched by diluting by about 10-fold with 8.0 M urea, 1 M Tris (2-chloroethyl) phosphate, pH 3.0. Sample preparation was performed using an HDx-3 PAL (LEAP Technologies). Data was collected at 30 s, 60 s, 120 s, 240 s, 480 s, 960 s, 1920 s, and 3840 s in three independent labeling experiments. After quenching, the solutions were subjected to online pepsin digestion followed by LC/MS analysis using UltiMate3000RSLCnano (Thermo Fisher Scientific) connected to a Q Exactive(tm) Plus mass spectrometer (Thermo Fisher Scientific). Online pepsin digestion was performed with a Poroszyme® Immobilized Pepsin Cartridge 2.1 × 30 mm (Waters Corporation) in formic acid solution (pH 2.5) for 3 min at 8 °C, at a flow rate of 50 µL min^-1^. Desalting and analytical processes after pepsin digestion were performed using Acclaim(tm) PepMap(tm) 300 C18 (1.0 × 15 mm; Thermo Fisher Scientific) and Hypersil GOLD(tm) (1.0 × 50 mm; Thermo Fisher Scientific) columns. The mobile phase was 0.1% formic acid solution (pH 2.5) (buffer A) and 0.1% formic acid (pH 2.5) containing 90% acetonitrile (B buffer). The deuterated peptides were eluted at a flow rate of 45 μL min^-1^ with a gradient of 10% – 90% of buffer B for 9 min. The conditions of the mass spectrometer were as follows: electrospray voltage, 3.8 kV; positive ion mode; sheath and auxiliary nitrogen flow rates at 20 and 2 arbitrary units, respectively; ion transfer tube temperature at 275 °C; auxiliary gas heater temperature at 100 °C; and a mass range of m/z 200 – 2,000. Data-dependent acquisition was performed using a normalized collision energy of 27 arbitrary units. Analysis of the deuteration levels of the peptide fragments was performed based on the MS raw files, by comparing the spectra of deuterated samples with those of non-deuterated samples, using the HDExaminer software (Sierra Analytics).

## Supporting information

This article contains supporting information.

## Data availability statement

All data are available from the authors upon request. Please send request to Satoru Nagatoishi.

## Acknowledgments and Funding Information

We thank Thermo Fisher Scientific for the technical support in HDX-MS experiments. This work was supported by JSPS KAENHI under grant number JP18H02082, JP18H05425 (to S.N.) and JP16H02420, JP19H05766, JP20H02531 (to K.T.).

## Footnotes

This article contains supporting information.

## Abbreviations

IL-11: Interleukin-11
IL-11Rα: lL-11 receptor alpha
IL-6: Interleukin-6
IL-6Rα: lL-6 receptor alpha
pg130: glycoprotein 130
SPR: surface plasmone resonance
ITC: isothermal titration calorimetry
HDX-MS: hydrogen/deuterium exchange mass spectrometry
BSA: Buried surface area

