## Supporting Information for "Biophysical insight into protein-protein interactions in the Interleukin-11/Interleukin-11Rα/glycoprotein 130 signaling complex"

**Running title:**

Biophysical analysis of IL-11 signaling complex

**Expression and purification of interleukins**

The sequence of human IL-11 (amino acids 23-199) containing an N-terminal hexahistidine-tag was subcloned into a pET28b vector. *Escherichia coli* BL21 (DE3) competent cells (EMD Millipore) were transformed with the plasmids and grown at 37 °C in LB medium with 50 μL $ml^{-1}$ ampicillin. The cells were cultured to $OD_{600}$ = 0.6. Protein expression was induced by the addition of 0.5 μM IPTG and further cultured at 20 °C for 16 h. The cells were harvested by centrifugation at 8,000 × *g* for 15 min at 4 °C. After sonication of the cells, cell lysates were cleared by centrifugation, and IL-11 was purified from the lysates using Ni-NTA agarose (Qiagen) at 4 °C. Bound proteins were eluted with 20 mM Tris-HCl (pH 8.0), 500 mM NaCl, 500 mM imidazole. Final purifications were performed using SEC on a HiLoad® 26/600 Superdex® 75 pg (GE Healthcare) with running buffer composed of 20 mM HEPES-NaOH (pH 7.4) and 150 mM NaCl.

The sequence of human IL-6 (amino acids 30-212) containing a myc-tag and hexahistidine-tag at C-terminus was subcloned into pcDNA™ 3.4-TOPO® vector (Thermo Fisher Scientific). The vector was transfected into Expi293F™ cells (Thermo Fisher Scientific) according to the manufacturer's protocol. Enhancer 1 and Enhancer 2 were added to transfected cultures at 20 h post-transfection. The cells were cultured at 37 °C, 125 rpm, 7% $CO_2$ for 3 days. IL-6 was purified from the supernatant using Ni-NTA agarose (Qiagen) at 4 °C. Bound protein was eluted with 20 mM Tris-HCl (pH 8.0)

500 mM NaCl 500 mM imidazole. Final purifications were performed by SEC on HiLoad® 16/600 Superdex® 200 pg (GE Healthcare) with running buffer composed of 20 mM HEPES-NaOH (pH 7.4) and 150 mM NaCl.

**Expression and purification of interleukin receptors**

The gene encoding the extracellular domain of IL-11Rα (amino acids 23-363) containing a sequence of the Fc domain of human IgG and FLAG-tag at the C-terminus was subcloned into a pFastBac1™ vector (Invitrogen). In addition, the sequence encoding an SP1-2 signal peptide described previously (28) was added at the N-terminus. Recombinant baculovirus was obtained using the Bac-to-Bac Baculovirus expression system (Invitrogen). Mimic™ Sf9 Insect Cells (Gibco™) at a cell density of $1.8 \times 10^8$ cells・$mL^{-1}$ were infected with P3 virus stock solution at 1% (v/v). The cells were cultured in Sf-900™ II SFM (Gibco™) supplemented with 10% (v/v) fetal bovine serum at 27 °C, 125 rpm for 4 days. IL-11Rα was purified from the supernatant by using anti-FLAG M2 affinity gel (Sigma-Aldrich) at 4 °C according to the manufacturer's protocol. Bound protein was eluted with 1 M arginine-HCl (pH 4.4), and the column eluate was neutralized with 2 M Tris-HCl (pH 8.0). Final purification was performed by SEC on HiLoad® 16/600 Superdex® 200 pg (GE Healthcare) with running buffer composed of 20 mM HEPES-NaOH (pH 7.4) and 150 mM NaCl.

The sequence of human IL-6Rα (amino acids 20-358) containing a sequence of Fc domain of human IgG, a myc-tag, and a hexahistidine-tag at the C-terminus was subcloned into plasmid pcDNA™ 3.4-TOPO® vector (Thermo Fisher Scientific). The sequence of human gp130 (24-618) containing a myc-tag and a hexahistidine-tag at the C-terminus was subcloned into pcDNA™ 3.4-TOPO® vector (Thermo Fisher Scientific). For the expression of IL-6Rα and gp130, the Expi293™ Expression System (Thermo Fisher Scientific) was used. The transfection of Expi293F™ cells was performed according to the manufacturer's protocol. Enhancer 1 and Enhancer 2 were added to transfected cultures at 20 h post-transfection. The cells were cultured at 37 °C, 125 rpm, 7% $CO_2$ for 3 days. Protein was purified from the supernatant by using Ni-NTA agarose (QIAGEN) at 4 °C. Bound proteins were eluted with 20 mM Tris-HCl (pH 8.0) 500 mM NaCl 500 mM imidazole. Final purifications were performed by SEC on HiLoad® 16/600 Superdex® 200 pg (GE Healthcare) with running buffer composed of 20 mM HEPES-NaOH (pH 7.4) and 150 mM NaCl.

**Table S1. Kinetic parameters of the interaction between IL-11 and IL-11Rα**

| | $k_{on}$ ($M^{-1}s^{-1}$) | $k_{off}$ ($s^{-1}$) | $K_D$ (nM) |
|---|---|---|---|
| 10 ºC | $1.2 \times 10^6$ | $1.7 \times 10^{-3}$ | 1.5 |
| 15 ºC | $1.2 \times 10^6$ | $1.4 \times 10^{-3}$ | 1.2 |
| 20 ºC | $4.3 \times 10^6$ | $3.0 \times 10^{-3}$ | 0.69 |
| 25 ºC | $6.1 \times 10^6$ | $3.2 \times 10^{-3}$ | 0.53 |
| 30 ºC | $1.3 \times 10^7$ | $5.4 \times 10^{-3}$ | 0.43 |

**Table S2. Kinetic parameters of the interaction between IL-6 and IL-6Rα**

| | $k_{on}$ ($M^{-1}s^{-1}$) | $k_{off}$ ($s^{-1}$) | $K_D$ (nM) |
|---|---|---|---|
| 10 ºC | $3.2 \times 10^5$ | $2.8 \times 10^{-3}$ | 8.9 |
| 13 ºC | $1.1 \times 10^6$ | $1.1 \times 10^{-2}$ | 9.8 |
| 15 ºC | $6.9 \times 10^5$ | $7.9 \times 10^{-3}$ | 12 |
| 20 ºC | $1.7 \times 10^6$ | $3.1 \times 10^{-2}$ | 18 |

**Table S3. Thermodynamic parameters of the interactions between interleukins and their receptors based on SPR data**

| | $\Delta H$ (kJ/mol) | $-T\Delta S$ (kJ/mol) | $\Delta G$ (kJ/mol) |
|---|---|---|---|
| IL-11 with IL-11Rα | 47 ± 3 | −100 ± 3 | −53 ± 4 |
| IL-6 with IL-6Rα | −49 ± 5 | 6 ± 5 | −43 ± 7 |

**Table S4. Thermodynamic parameters of the interactions between gp130 and the IL-6/IL-6Rα and IL-11/IL-11Rα based on ITC data**

| | $\Delta H$ (kJ/mol) | $-T\Delta S$ (kJ/mol) | $\Delta G$ (kJ/mol) | $K_D$ (nM) | Binding stoichiometry |
|---|---|---|---|---|---|
| gp130 with IL-11/IL-11Rα | −51 ± 1 | 5 ± 1 | −47 ± 1 | 6.9 ± 2.4 | 1.0 |
| gp130 with IL-6/IL-6Rα | −28 ± 1 | −13 ± 2 | −42 ± 1 | 54 ± 24 | 1.2 |

**Table S5A. Buried surface areas (BSA) of amino acids between IL-11 and gp130 D2 in interleukin signaling complexes[a]**

| IL-11 | BSA (Å) | gp130 D2 | BSA (Å) |
|---|---|---|---|
| Leu 243 | 32.91 | Trp 141 | 33.14 |
| Ser 246 | 39.47 | Thr 143 | 78.31 |
| Arg 330 | 53.62 | Ser 165 | 0.0 |
| Arg 333 | 150.47 | Val 167 | 47.47 |
| Arg 337 | 49.87 | Tyr 168 | 0.0 |
| | | Phe 169 | 107.76 |
| | | Val 170 | 57.17 |

[a] Data for the IL-11 complex is from the predicted structure. A blank cell indicates no difference between value for gp130 alone and for gp130 in the complex.

**Table S5B. Buried surface areas (BSA) of amino acids between IL-6 and gp130 D2 in the interleukin signaling complex[a]**

| IL-6 | BSA (Å) | gp130 D2 | BSA (Å) |
|---|---|---|---|
| Gln 28 | 34.66 | Trp 141 | 70.02 |
| Tyr 31 | 87.87 | Thr 143 | 0.0 |
| Glu 110 | 76.79 | Ser 165 | 78.67 |
| Ala 1145 | 15.05 | Val 167 | 42.83 |
| Met 117 | 63.99 | Tyr 168 | 32.27 |
| Ser 118 | 12.65 | Phe 169 | 127.45 |
| Val 121 | 45.18 | Val 170 | 16.36 |
| Gln 124 | 53.49 | | |
| Phe 125 | 31.72 | | |

[a] Data for the IL-6 complex are from the crystal structure, PDB ID: 1p9m. A blank cell indicates no difference between value for gp130 alone and for gp130 in the complex.

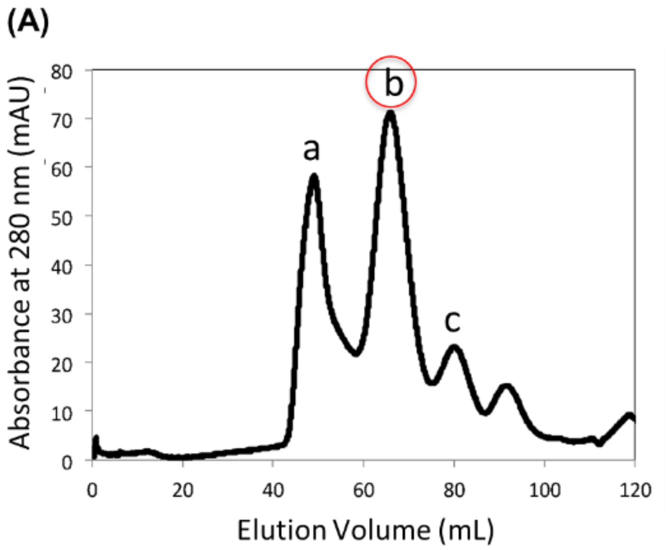


**Figure S1. Preparation of IL-11Rα.** (A) SEC sensorgram of IL-11Rα in the purification process. (B) SDS-PAGE of fractions from SEC. Fraction b was used in experiments.

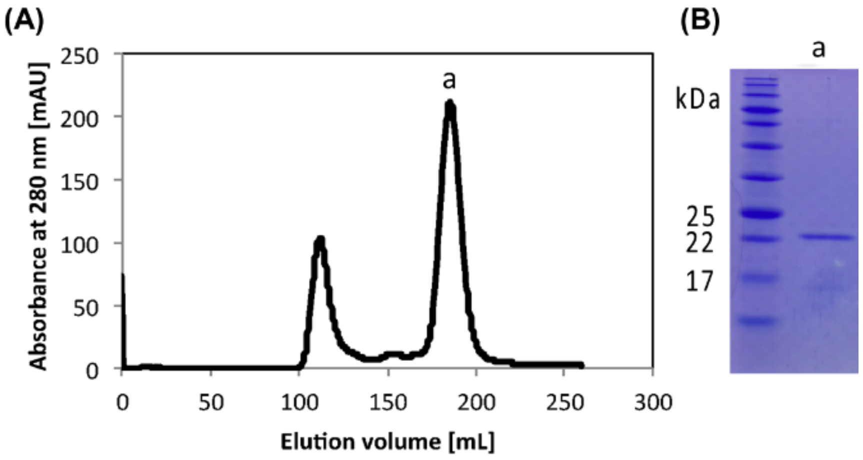


**Figure S2. Preparation of IL-11.** (A) SEC sensorgram of IL-11 in the purification process. (B) SDS-PAGE of SEC fractions. Fraction a was used in experiments.

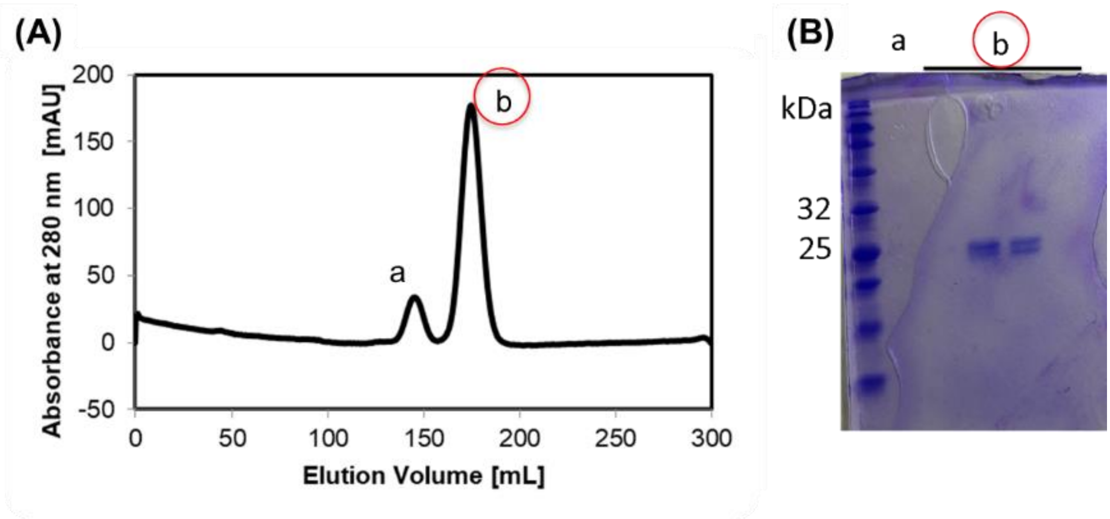


**Figure S3. Preparation of IL-6.** (A) SEC sensorgram of IL-6 in the purification process. (B) SDS-PAGE of SEC fractions. Fraction b was used in experiments.

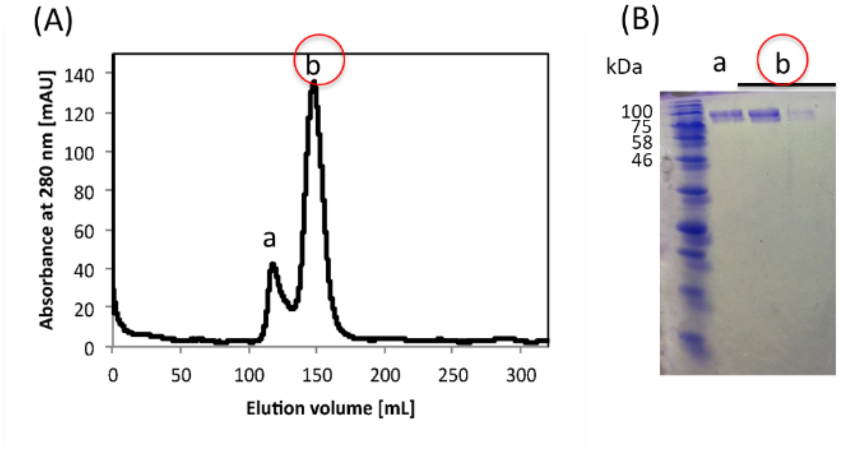


**Figure S4. Preparation of IL-6Rα.** (A) SEC sensorgram of IL-6 Rα in the purification process. (B) SDS-PAGE of SEC fractions. Fraction b was used in experiments.

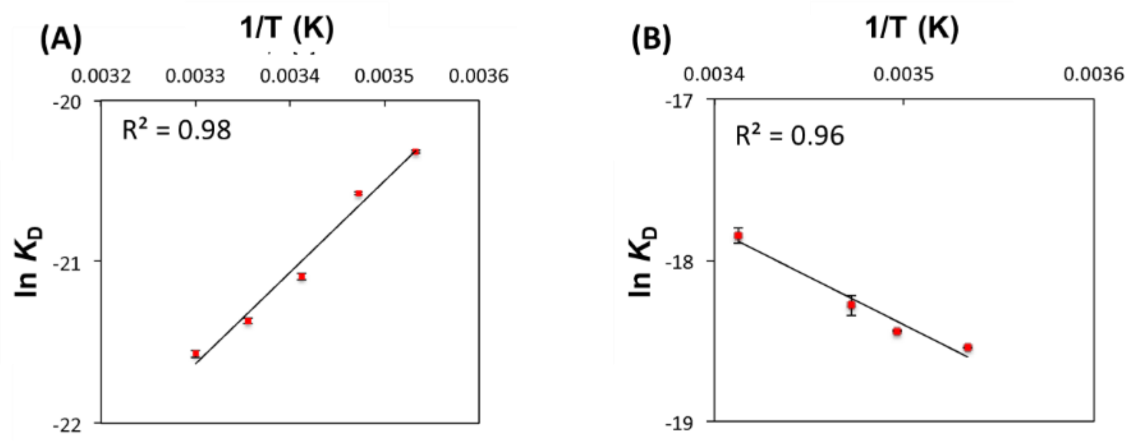


**Figure S5. van't Hoff thermodynamic analysis of the interaction between (A) IL-11 and IL-11Rα and (B) IL-6 and IL-6Rα.**

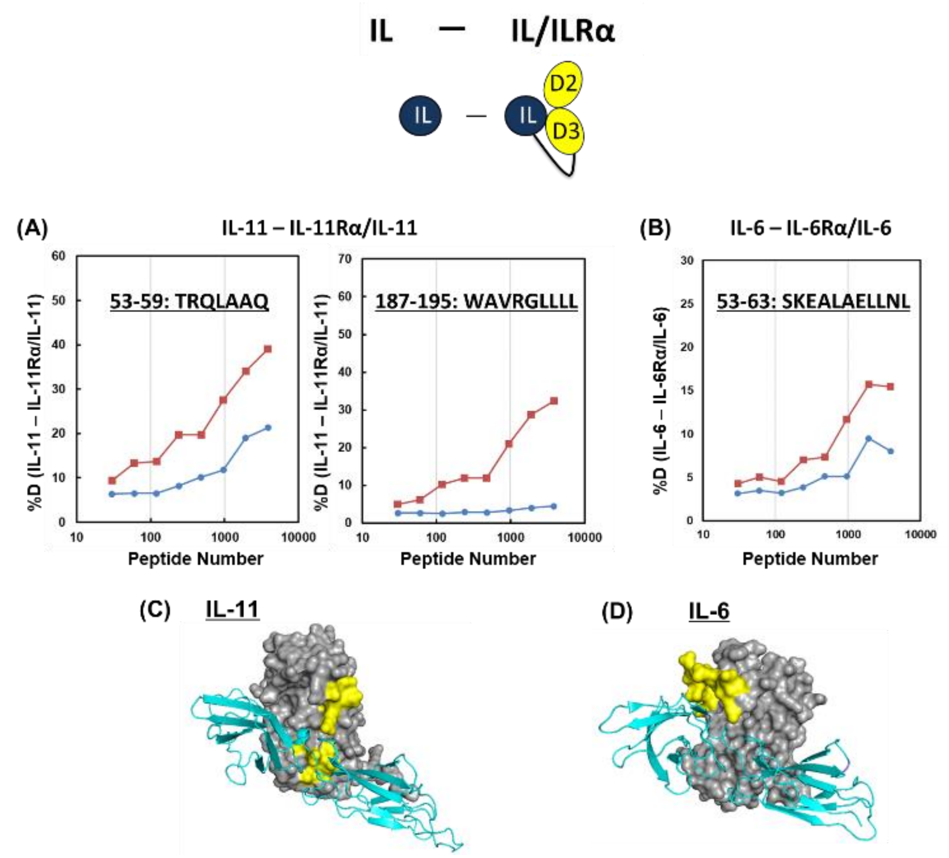


**Figure S6. HDX-MS analyses of interleukin/receptor complexes.** (A) HDX-MS profiles of indicate regions of IL-11 in the absence (red line) and presence (blue line) of IL-11Rα. (B) HDX-MS profiles of indicated regions of IL-6 in the absence (red line) and presence (blue line) of IL-6Rα. (C, D) Models of C) IL-11/IL-11Rα and D) IL-6/IL-6Rα. The yellow color represents regions where deuterium exchange ratio decreased in the complex. The interleukin is shown as a gray surface model, and the receptor is in cyan.

### (A) gp130 D1 (41-98) and IL-11 contact

| | | BSA (Å$^2$) |
|---|---|---|
| ASN | 75 | 0.00 |
| ILE | 76 | 0.00 |
| GLN | 77 | 59.67 |
| LEU | 78 | 0.00 |
| THR | 79 | 24.99 |
| CYS | 80 | 0.00 |
| ASN | 81 | 5.96 |
| ILE | 82 | 0.00 |
| LEU | 83 | 0.00 |
| THR | 84 | 0.00 |
| PHE | 85 | 0.00 |
| GLY | 86 | 0.00 |
| GLN | 87 | 0.00 |
| LEU | 88 | 0.00 |
| GLU | 89 | 31.72 |
| GLN | 90 | 57.51 |
| ASN | 91 | 74.48 |
| VAL | 92 | 9.37 |
| TYR | 93 | 40.47 |
| GLY | 94 | 34.53 |
| ILE | 95 | 0.67 |
| THR | 96 | 17.31 |
| ILE | 97 | 0.00 |
| ILE | 98 | 0.00 |

### (B) gp130 D1 (41-98) and IL-6 contact

| | | BSA (Å$^2$) |
|---|---|---|
| ASN | 75 | 0.00 |
| ILE | 76 | 0.00 |
| GLN | 77 | 20.24 |
| LEU | 78 | 0.00 |
| THR | 79 | 11.72 |
| CYS | 80 | 0.00 |
| ASN | 81 | 0.00 |
| ILE | 82 | 3.97 |
| LEU | 83 | 0.00 |
| THR | 84 | 0.00 |
| PHE | 85 | 0.00 |
| GLY | 86 | 0.00 |
| GLN | 87 | 0.00 |
| LEU | 88 | 0.00 |
| GLU | 89 | 29.98 |
| GLN | 90 | 52.67 |
| ASN | 91 | 32.65 |
| VAL | 92 | 8.63 |
| TYR | 93 | [illegible]3.00 |
| GLY | 94 | 28.52 |
| ILE | 95 | 5.73 |
| THR | 96 | 5.42 |
| ILE | 97 | 0.00 |
| ILE | 98 | 0.00 |

### (C) gp130 D3 and IL-11Rα contact

| | | BSA (Å$^2$) |
|---|---|---|
| SER | 247 | 0.00 |
| GLN | 248 | 0.00 |
| ILE | 249 | 0.00 |
| PRO | 250 | 0.00 |
| PRO | 251 | 0.00 |
| GLU | 252 | 17.24 |
| ASP | 253 | 45.07 |
| THR | 254 | 0.00 |
| ALA | 255 | 0.00 |
| SER | 256 | 28.38 |
| THR | 257 | 35.86 |
| ARG | 258 | 69.63 |
| SER | 259 | 24.52 |
| SER | 260 | 37.30 |
| PHE | 261 | 26.46 |
| THR | 262 | 41.81 |
| VAL | 263 | 0.00 |
| GLN | 264 | 94.48 |
| ASP | 265 | 36.49 |
| LEU | 266 | 0.00 |
| LYS | 267 | 0.00 |
| PRO | 268 | 0.00 |
| PHE | 269 | 0.00 |
| THR | 270 | 0.00 |

### (D) gp130 D3 and IL-6Rα contact

| | | BSA (Å$^2$) |
|---|---|---|
| SER | 247 | 0.00 |
| GLN | 248 | 0.00 |
| ILE | 249 | 0.00 |
| PRO | 250 | 0.00 |
| PRO | 251 | 0.00 |
| GLU | 252 | 28.29 |
| ASP | 253 | 50.50 |
| THR | 254 | 0.00 |
| ALA | 255 | 0.00 |
| SER | 256 | 41.98 |
| THR | 257 | 21.44 |
| ARG | 258 | 74.56 |
| SER | 259 | 6.35 |
| SER | 260 | 18.57 |
| PHE | 261 | 19.32 |
| THR | 262 | 49.60 |
| VAL | 263 | 0.16 |
| GLN | 264 | 45.71 |
| ASP | 265 | 4.42 |
| LEU | 266 | 0.00 |
| LYS | 267 | 0.00 |
| PRO | 268 | 0.00 |
| PHE | 269 | 0.00 |
| THR | 270 | 0.00 |

**Figure S7. Buried surface areas (BSA) of amino acids of gp130 D1 (41-98) and D3 in interleukin signaling complexes.**

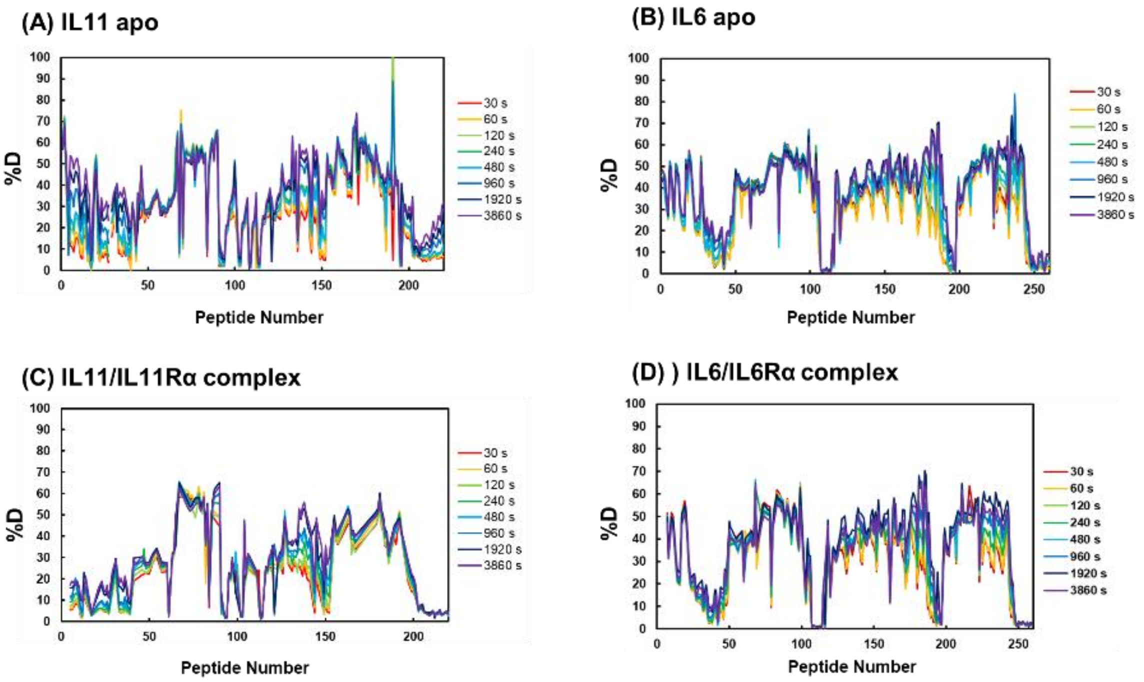


**Figure S8. HDX-MS of interleukins in the absence and presence of the receptor.**

HDX-MS profiles of (A) IL-11, (B) IL-6, (C) IL-11/IL-11Rα, and (D) IL-6/IL-6Rα.
